# Genetic, inflammatory, and epithelial cell differentiation factors control expression of human calpain-14

**DOI:** 10.1101/359638

**Authors:** Daniel E. Miller, Carmy Forney, Mark Rochman, Stacey Cranert, Jeffery Habel, Jeffrey Rymer, Arthur Lynch, Connor Schroeder, Josh Lee, Amber Sauder, Quinton Smith, Mehak Chawla, Michael P. Trimarchi, Xiaoming Lu, Ellen Fjellman, Michael Brusilovsky, Artem Barski, Stephen Waggoner, Matthew T. Weirauch, Marc E. Rothenberg, Leah C. Kottyan

## Abstract

Eosinophilic esophagitis (EoE) is a chronic, food-driven allergic disease resulting in eosinophilic esophageal inflammation. We recently found that EoE susceptibility is associated with genetic variants in the promoter of *CAPN14*, a gene with reported esophagus-specific expression. *CAPN14* is dynamically up-regulated as a function of EoE disease activity and after exposure of epithelial cells to interleukin-13 (IL-13). Herein, we aimed to explore molecular modulation of *CAPN14* expression. We identified three putative binding sites for the IL-13-activated transcription factor STAT6 in the promoter and first intron of *CAPN14*. Luciferase reporter assays revealed that the two most distal STAT6 elements were required for the ~10-fold increase in promoter activity subsequent to stimulation with IL-13 or IL-4, and also for the genotype-dependent reduction in IL-13-induced promoter activity. One of the STAT6 elements in the promoter was necessary for IL-13-mediated induction of *CAPN14* promoter activity while the other STAT6 promoter element was necessary for full induction. Chromatin immunoprecipitation in IL-13 stimulated esophageal epithelial cells was used to further support STAT6 binding to the promoter of *CAPN14* at these STAT6 binding sites. The highest *CAPN14* and calpain-14 expression occurred with IL-13 or IL-4 stimulation of esophageal epithelial cells under culture conditions that allow the cells to differentiate into a stratified epithelium. This work corroborates a candidate molecular mechanism for EoE disease etiology in which the risk variant at 2p23 dampens mediated *CAPN14* expression in differentiated esophageal epithelial cells following IL-13/STAT6 induction of *CAPN14* promoter activity.

## INTRODUCTION

Eosinophilic esophagitis (EoE) is a chronic, allergic disease associated with marked mucosal eosinophil accumulation (Liacouras et al. 2011). EoE remits after removal of specific food types, and food re-introduction causes disease reoccurrence, including dysregulation of esophageal transcripts. The etiology of EoE includes environmental, immunological, and genetic components (Alexander et al. 2014; Kottyan et al. 2014; Kottyan et al. 2015). One of the central questions in the EoE field, and allergy in general, is to understand why individuals develop certain manifestations of disease.

Until recently, studies have identified EoE-genetic risk loci that were broadly linked to other allergic diseases (Sherrill and Rothenberg 2014). For example, genetic variants at 5q22, encoding *TSLP* and *WDR36*, and 11q13, encoding *LRRC32* and *EMSY*, have been associated with allergic sensitization, asthma, allergic rhinitis, atopic dermatitis, food allergy and/or EoE (Greisenegger et al. 2013; Tang et al. 2012; Ferreira et al. 2011; Li et al. 2015; Ferreira et al. 2014; Weidinger et al. 2013; Ramasamy et al. 2011; Hinds et al. 2013; Sleiman et al. 2014). The shared association of risk variants among these phenotypes suggests that these loci contain variants that participate in the allele-dependent regulation of a molecular pathway that is central to the etiology of allergic disease.

We recently found that, in addition to genetic risk loci for allergic sensitization, EoE susceptibility is linked to one or more genetic factors at 2p23, encoding the *CAPN14* gene (Kottyan et al. 2014). This genetic linkage has been replicated at genome-wide significance in multiple cohorts, as well as in a recent independent study (Sleiman et al. 2014), adding credence to the importance of the 2p23 genetic linkage. *CAPN14* (encoding calpain-14) belongs to the classical calpain sub-family, a set of calcium activated intracellular regulatory proteases (Sorimachi et al. 2011). In our initial studies, we identified *CAPN14* as dynamically up-regulated as a function of EoE disease activity as well as after exposure of epithelial cells to IL-13 (Davis et al. 2016). Expression quantitative trait loci (eQTL) analysis revealed that patients with active EoE expressed *CAPN14* in a genotype-dependent manner (Kottyan et al. 2014). While *CAPN14* was expressed at a higher level in individuals with EoE compared to those without EoE, patients with the risk genotype had decreased expression of *CAPN14* compared to patients with the non-risk genotype. Consistent with these findings, overexpression of the calpain-14 protein in esophageal epithelial cells leads to morphological changes and barrier defects independently of IL-13-mediated inflammation, while *CAPN14* gene silencing in these cells leads to defects in barrier repair after IL-13 stimulation (Davis et al. 2016), suggesting that calpain-14 might contribute to EoE via a regulatory loop (Litosh et al. 2017).

In this study, we aimed to identify regulatory factors that control the expression of *CAPN14*. Using luciferase reporter constructs, we determined that IL-13 is sufficient to induce *CAPN14* promoter activity and identified one putative STAT6 binding sites as necessary for IL-13-induced *CAPN14* promoter activity with a second putative STAT6 binding site in the promoter necessary for full induction of *CAPN14* promoter activity. Using chromatin immunoprecipitation, we identified STAT6 binding to the promoter of *CAPN14* at the site of the two putative STAT6 binding sites in EPC2 esophageal epithelial cells. One single nucleotide polymorphism (SNP) that is highly associated with EoE risk and located in the promoter of *CAPN14* was sufficient to decrease *CAPN14* promoter activity in a manner that is consistent with the eQTL seen in individuals with and without EoE (Kottyan et al. 2014). By measuring mRNA and protein expression of CAPN14/calpain-14 over a variety of *in vitro* culture conditions, we further found that *CAPN14* expression is highest in differentiated esophageal epithelial cells after IL-13 exposure. IL-4 also signals in esophageal epithelial cells in a STAT6-dependent manner, and we obtained consistent results for IL-4 and IL-13 promoter activity and gene expression. Altogether, this study identified immunological, genetic, and epithelial cell differentiation mechanisms that regulate the expression of *CAPN14*.

## METHODS

### Esophageal epithelial cell cultures

EPC2 esophageal epithelial cells were grown in keratinocyte serum-free media (K-SFM) (Life Technologies, Grand Island, NY) to various levels of confluence (80% or 100%) with relatively low (0.09 mM) or high (1.8 mM) Ca^2+^ and with or without 100 ng/mL IL-13 or IL-4 for 24 hours, as a monolayer submerged culture or in an air-liquid interface (ALI) culture (Davis et al. 2016; Kottyan et al. 2014). For the ALI culture system, the esophageal epithelial EPC2 cell line was grown to confluence on 0.4 μm pore-size polyester permeable supports (Corning Incorporated, Corning, NY) in K-SFM supplemented with 1.8 mM calcium. Epithelial differentiation was then induced by removing culture media from the inner chamber of the permeable support and maintaining the esophageal epithelial cells for 8 days in the ALI in the presence or absence of IL-13 or IL-4 (100 ng/mL). TE7 esophageal epithelial cells were grown in RPMI1640 supplemented with 10% FBS (HyClone, Pittsburgh, PA).

### Luciferase reporter assays

The *CCL26* nanoluciferase reporter construct was engineered using the firefly reporter construct reported previously (Lim et al. 2011). The first 964 bp before the *CCL26* start codon were subcloned into the pNL1.1 nanoluciferase reporter vector (Promega, Madison, WI). For the *CAPN14* promoter reporter construct, PCR primers (listed in **Table S1**) were used to amplify the 1988 base pair region around rs76562819 and subsequently clone it into the pNL1.1 nanoluciferase reporter vector. Site-directed mutagenesis was performed to generate the construct with scrambled STAT6 binding sites and the EoE non-risk allele of rs76562819 using the GeneArt Site-Directed Mutagenesis System (Invitrogen, Darmstadt, Germany). Sequences were confirmed by Sanger sequencing. All mutagenesis primers are provided in **Table S1**.

250 ng of nanoluciferase experimental constructs were transiently co-transfected into esophageal epithelial cells with 250 ng of pGL3-control firefly luciferase reporter plasmid. After 24 hours, cells were treated with 100 ng/mL of IL-13 or IL-4 (Promega, Madison, WI). 24 hours after cytokine administration, cells were lysed and nano and firefly luciferase activity was assayed using the Promega Nano-Glo Luciferase Assay System. The nanoluciferase measurement for each well was divided by the firefly luciferase measurement in order to account for cytotoxicity, small differences in cell number, and transfection efficiency. Each well was further normalized to the promoterless pNL1.1 vector to control for baseline activity of the vector.

### Chromatin immunoprecipitation analysis

EPC2 cells were treated with IL-13 at 100 ng/mL for 45 min and fixed with 1% formaldehyde for ChIP. ChIP was performed as reported previously (Rochman et al. 2015) with the exception that the sonication was for 4 min and ChIP was performed manually for the anti-STAT6 experiments. Anti-STAT6 antibody S-20 (D3H4 Rabbit mAb #5397, Cell Signaling) anti-H3K4me3 antibody ab8580 (Abcam), and anti-H3K27Ac antibody ab4279 (Abcam). We fixed 10–20×10^6^ cells with 0.8% formaldehyde by adding 1 ml of 10X fixation buffer (50 mM Hepes-KOH, pH 7.5; 100 mM NaCl; 1 mM EDTA; 0.5 mM EGTA; 8% formaldehyde) to 9 ml of growth medium for 8 minutes at room temperature with shaking. The reaction was stopped by adding glycine to a final concentration of 125 mM for an additional 5 minutes. After washing with PBS, pellets were frozen at −80°C for at least overnight. Nuclei were prepared by re-suspending pellets in 1 ml of L1 buffer (50 mM Hepes-KOH, pH 7.5; 140 mM NaCl; 1 mM EDTA; 10% glycerol; 0.5% NP-40; 0.25% Triton X-100) and incubated at 4°C for 10 min. Nuclei were pelleted and re-suspended in 1 ml of L2 buffer (10 mM Tris-HCl, pH 8.0; 200 mM NaCl; 1 mM EDTA, pH 8.0; 0.5 mM EGTA, pH 8.0) and rotated for 10 min at room temperature. Nuclei were briefly washed with sonication buffer (Tris-EDTA [TE] buffer + 0.1% SDS) and re-suspended in 1 ml of sonication buffer. All buffers were supplemented with complete EDTA-free protease inhibitors (Roche Diagnostics Corporation Indianapolis, IN). Sonication was performed by a Covaris S220 focused ultrasonicator (Covaris, Inc. Woburn, MA) at 175W Peak Incident Power, 10% output, 200 bursts for 10 min in 12×12-mm, round-bottom glass tubes. Efficient DNA fragmentation was verified by agarose gel electrophoresis. Anti-histone mark ChIP was performed by SX-8G IP-Star^®^ Automated System (Diagenode Inc., Denville, NJ) in RIPA buffer (TE + 0.1% SDS, 1% Triton X-100, 150 mM NaCl, 0.1% sodium deoxycholate) following the protocol of the manufacturer with 2–4 μg of indicated antibodies. Libraries were sequenced on the Illumina HiSeq 2500, and ChIP-seq data were visualized and analyzed in Biowardrobe (Kartashov and Barski 2015; Vallabh et al. 2018).

### RNA purification and expression analyses

For the real-time PCR analysis of CAPN14 expression, total RNA was purified from esophageal epithelial cells using the MirVana miRNA Isolation kit (ThermoFisher, Waltham, MA). RNA was reverse-transcribed with the High-Capacity RNA-to-cDNA kit (Applied Biosystems, Grand Island, NY). Gene expression was determined by real-time PCR using a 7500 Real-time PCR system (Applied Biosystems, Foster City, CA) using Taqman Gene Expression Assays CAPN14-Hs00871882_m1, CCL26- Hs00171146_m1 and GAPDH-Hs02786624_g1. Relative quantification was calculated by the comparative CT method (Livak and Schmittgen 2001). Briefly, expression levels for *CAPN14* or *CCL26* were normalized to levels of *GAPDH*. All samples were then normalized to cells grown at 80% confluence without high calcium or IL-13 and the data were exponentially transformed.

For the RNA-sequencing study performed to identify the isoforms of CAPN14 expressed in the esophageal biopsies of people with and without EoE (**Figure S2**), RNA was isolated from distal esophageal biopsy RNA from EoE patients (5 male and 5 female) with active disease and from unaffected controls (5 male and 5 female) as previously described (Kottyan et al. 2014; Blanchard et al. 2006; Namjou et al. 2014). EoE biopsies showed active disease pathology at the time when they were taken, and all patients reported no glucocorticoid treatment at the time of biopsy.

RNA sequencing acquiring 50 million mappable 125 base-pair reads from paired-end libraries was performed at the Genetic Variation and Gene Discovery Core Facility at CCHMC. Data were aligned to the GrCh37 build of the human genome using the Ensembl (Flicek et al. 2013) annotations as a guide for TopHat (Trapnell et al. 2009). Expression analysis was performed using Kallisto’s quant function (Bray et al. 2016). Data has been deposited in NCBI Geo database as GSE113341.

### Protein isolation and analysis

Whole cell lysate was extracted from EPC2 cells lysed with RIPA Buffer (Millipore, Burlington, MA) and Halt^™^ phosphatase inhibitor cocktail. Samples were exposed to electrophoresis in NuPAGE Novex 4-12% Bis-Tris Gels and transferred to nitrocellulose using the iBlot system (Invitrogen, Carlsbad, CA). Primary antibodies: anti-CAPN14 (HPA035721, Sigma Aldrich, St. Louis, MO) and anti-β-actin (4967L, Cell Signaling Technologies, Danvers, MA). Secondary antibody: IRDye 800CW donkey anti-rabbit (926-32213, LI-COR, Lincoln, NE). Nitrocellulose membranes were imaged using the Odyssey CLx system (LI-COR) and analyzed with Image Studio (LI-COR). EPC2 cells that constitutively express *CAPN14* behind a CMV promoter were used as a positive control.

Statistical analysis and data visualization were done with GraphPad Prism (GraphPad Software, La Jolla, CA).

### Data and reagent availability

Constructs are available upon request. For the *CAPN14* promoter reporter construct, PCR primers (listed in **Table S1**) were used to amplify the 1988 base pair region around rs76562819 and subsequently clone it into the pNL1.1 nanoluciferase reporter vector. All mutagenesis primers are provided in **Table S1**. RNA sequencing data has been deposited in NCBI Geo database as GSE113341.

## RESULTS

IL-13 administration to esophageal epithelial cells is sufficient to induce *CAPN14* expression similar to *CCL26*, a chemokine whose promoter activity is dependent on the STAT6 transcription factor (Lim et al. 2011). Based upon these previous results, we hypothesized that IL-13 acted through the transcription factor STAT6 to directly promote *CAPN14* expression. Thus, we first constructed promoter reporters by placing the promoters of either *CAPN14* or *CCL26* in front of the nanoluciferase gene (**Figure 1 A**). After treatment with IL-13 or IL-4, esophageal epithelial cells transfected with the *CAPN14* reporter construct exhibited a similar upregulation of nanoluciferase expression to the cells transfected with a *CCL26* reporter construct (**Figure 1 B and Figure S1 A**). Based upon the canonical 5’-TTC…GAA-3’ core motif of STAT6, we identified two putative STAT6 binding sequences in the promoter located within 90 base pairs of the transcription start site of *CAPN14* and one putative STAT6 binding site in the first intron of *CAPN14*. We tested the necessity of each individual predicted STAT6 binding site for IL-13-induced promoter activity by creating luciferase reporter constructs with mutated putative STAT6 sites (**Figure 1 A**). Mutation of the first STAT6 site attenuated IL-13-mediated induction of CAPN14 promoter activity, while mutation of the second STAT6 site completely blocked CAPN14 promoter induction (**Figure 1 C**). Mutation of the third putative STAT6 site in the first intron of CAPN14 did not affect the promoter activity of CAPN14 following treatment with IL-13 (**Figure 1 C**). From these experiments, we concluded that the two sites in the promoter are necessary for full IL-13-induced promoter activity.

**Figure 1.**
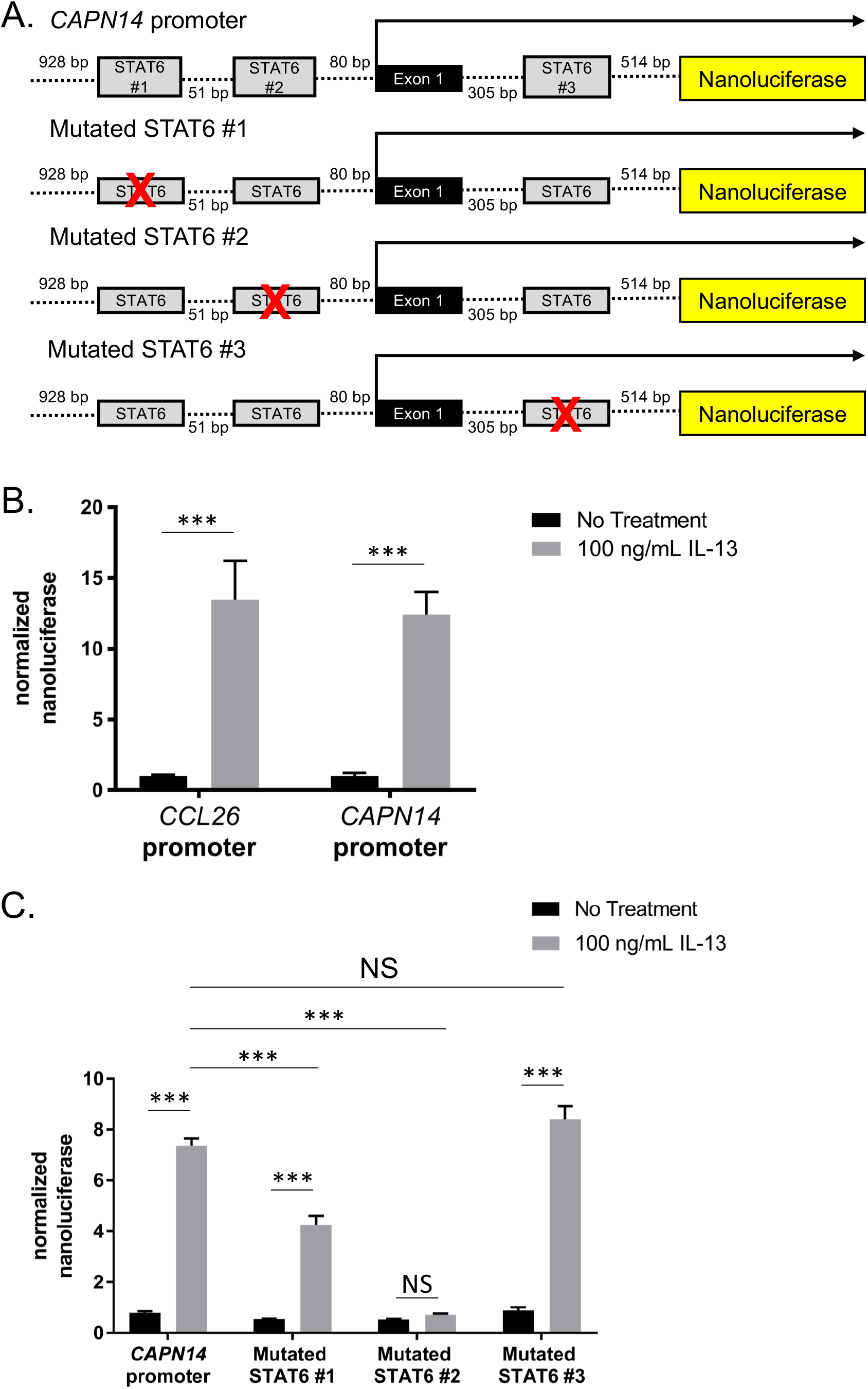
STAT6 binding sites in the promoter region of the *CAPN14* gene affect IL-13-induced *CAPN14* promoter activity. A, schematic representation of the nucleotide sequence of the promoter of the human *CAPN14* gene. Putative binding sites for STAT6 are defined by grey boxes. B, Site-directed mutagenesis elimination of putative STAT6-binding sites is denoted by crossed boxes. The (B) CCL26 and CAPN14 constructs or the (C) mutagenized CAPN14 constructs were transfected into esophageal epithelial cells followed by treatment with or without IL-13 (100 ng/mL) for 24 h. For each sample, nanoluciferase activity was normalized to firefly luciferase activity. Data are shown as mean ± S.E.M. (***, *p* < 0.001; *n* = 3 per group; data representative of three independent experiments).

Chromatin immunoprecipitation of STAT6 in esophageal epithelial cells treated with IL-13 was used to further establish STAT6 binding to the promoter of *CAPN14* at the two binding sites that were necessary for IL-13-inducible *CAPN14* promoter activity. We identified ChIP-seq peaks in the promoter of *CAPN14* spanning these two STAT6 sites from chromatin extracted from IL-13 stimulated esophageal epithelial cells (**Figure 2**). There was no evidence of STAT6 binding at the third putative STAT6 site in the first intron of *CAPN14*. These STAT6 ChIP-seq peaks overlap ChIP-seq peaks from active histone marks H3K4me3 and H3K27Ac that appear after induction by IL-13 (**Figure 2**). Altogether, these functional genomic data support our initial hypothesis that IL-13 and IL-4 activate *CAPN14* expression through the binding of STAT6 to two sites in the promoter of *CAPN14*.

**Figure 2.**
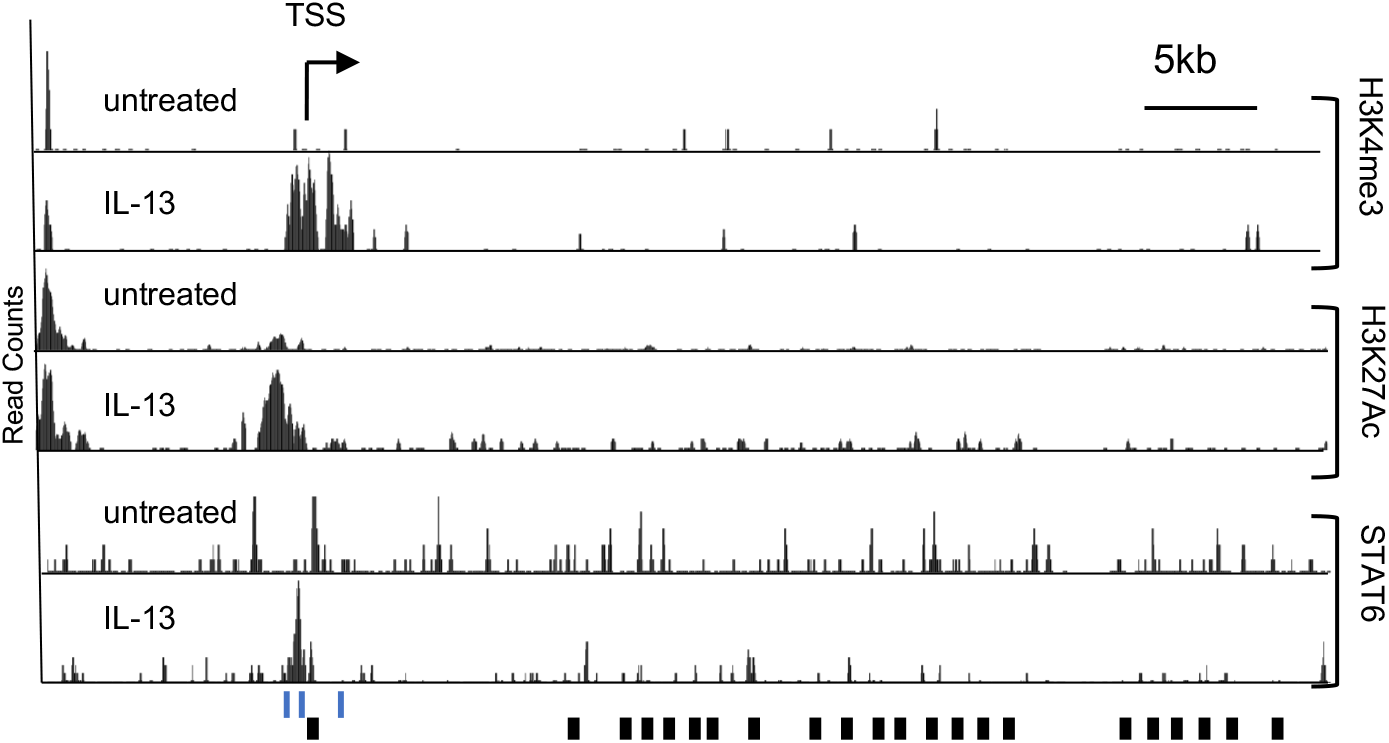
Epigenetic signature and STAT6 binding to the promoter region of *CAPN14* in response to IL-13 stimulation. An image capture from the UC Santa Cruz genome browser shows the tag density profile of aligned ChIP-seq reads in untreated and IL-13-treated EPC2 cells. The read depths of the aligned sequences are of the same scale. Arrow represents direction of transcription; black boxes represent exons. In the IL-13-treated cells, elevated levels of H3K4me3 and H3K27Ac are detected in the promoter region. The scale is the same for all track pairs (untreated and IL-13). Three putative STAT6 binding sites around the TSS are indicated by blue lines. TSS – transcription start site.

Patients with EoE and the genetic-risk haplotype at 2p23, marked by single nucleotide polymorphism (SNP) rs76562819, have 50% less *CAPN14* in their esophageal biopsies compared to patients with the non-risk haplotype (Kottyan et al. 2014). An analysis of differential isoform usage of *CAPN14* identified two major isoforms of CAPN14 that did not change with EoE disease status or sex, (**Figure S2**). Of the two isoforms of *CAPN14* that are expressed, the major isoform (ENST00000444918) includes exon 7, while the less highly expressed isoform of *CAPN14* (ENST00000398824) does not. Furthermore, the less highly expressed isoform of *CAPN14* (ENST00000398824) is predicted to undergo nonsense mediated decay based upon manual annotation from the VEGA/Havana projects (Wilming et al. 2008). The EoE-risk haplotype at 2p23 includes 12 variants in linkage disequilibrium (r^2^0.8), but functional genomic and biochemical evidence supported a specific role for rs76562819 (Davis et al. 2016; Kottyan et al. 2014). We tested the hypothesis that rs76562819 was sufficient to result in genotype-dependent promoter activity of *CAPN14* using two identical luciferase constructs containing the promoter and a portion of the first intron of *CAPN14* with either the risk or the non-risk allele of rs76562819 (**Figure 3 A**). We found that the EoE risk allele at rs76562819 resulted in a 40.0% reduction of IL-13 and IL-4-induced *CAPN14* promoter activity compared to the EoE non-risk allele (**Figure 3 B**). Thus, the single base change from non-risk to risk at rs76562819 is sufficient to explain most of the reduced *CAPN14* promoter activity in the luciferase reporter assays. These results are consistent with the genotype at rs76562819 accounting for genotype-dependent expression observed in EoE patient biopsies.

**Figure 3.**
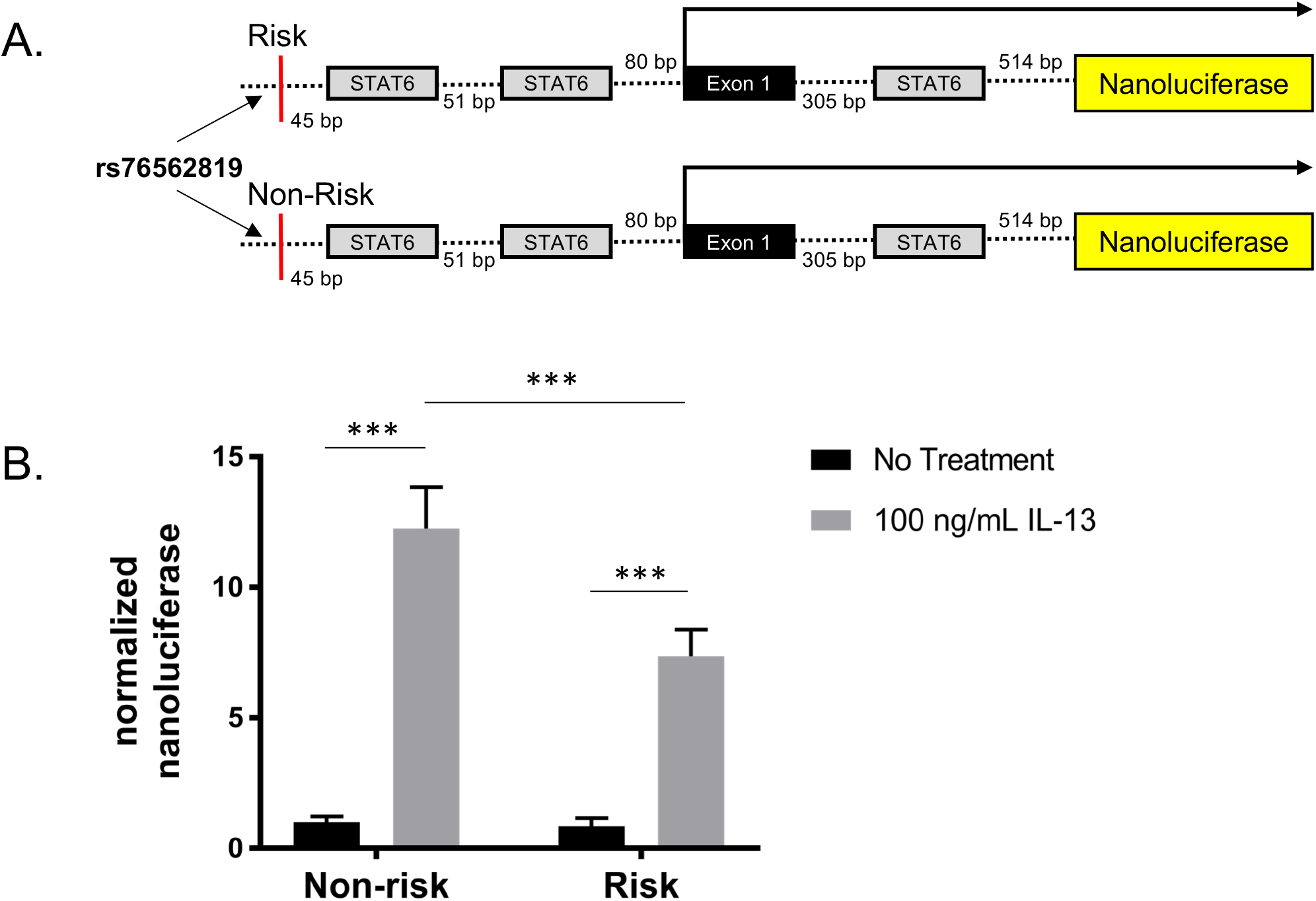
The risk variant of rs76562819 results in a genotype-dependent decrease in IL-13 induced promoter activity of *CAPN14*. *A*, Diagrams of constructs containing the *CAPN14* promoter are shown. *B*, Reporter constructs were transfected into esophageal epithelial cells followed by treatment with or without IL-13 (100 ng/mL) for 24 h. For each sample, nanoluciferase activity was normalized to firefly luciferase activity. Data are shown as mean ± S.E.M. (***, t-test p-value<0.001; 2-way ANOVA: p<0.05, genotype p<0.0001 accounting for 12.6% of total variation; *n* = 3 per group; data representative of three independent experiments).

We next sought to determine the factors leading to expression of *CAPN14* in esophageal epithelium. Unlike *CCL26, CAPN14* expression is highly dependent upon the cell culture conditions. Esophageal epithelial differentiation is a physiological process that increases mechanical strength and barrier function to squamous epithelia (Macara et al. 2014). Factors that are necessary for differentiation *in vitro* are calcium concentration, cellular confluence, and the culture system (Macara et al. 2014; Kc et al. 2015). Without esophageal epithelial differentiation by confluence, calcium, and growth in an air-liquid interface (ALI), *CAPN14* mRNA expression is low (**Figure 4 A**). Following differentiation, *CAPN14* is induced 10,000-fold in the ALI, p<0.0001 (**Figure 4 A**). At a protein level, calpain-14 was detected only in the most differentiated system (ALI) following treatment with IL-13 (**Figure 4 B**). As seen in the promoter reporter experiments, IL-4 and IL-13 are equally capable of induction of *CAPN14* mRNA and protein (**Figure S3**).

**Figure 4.**
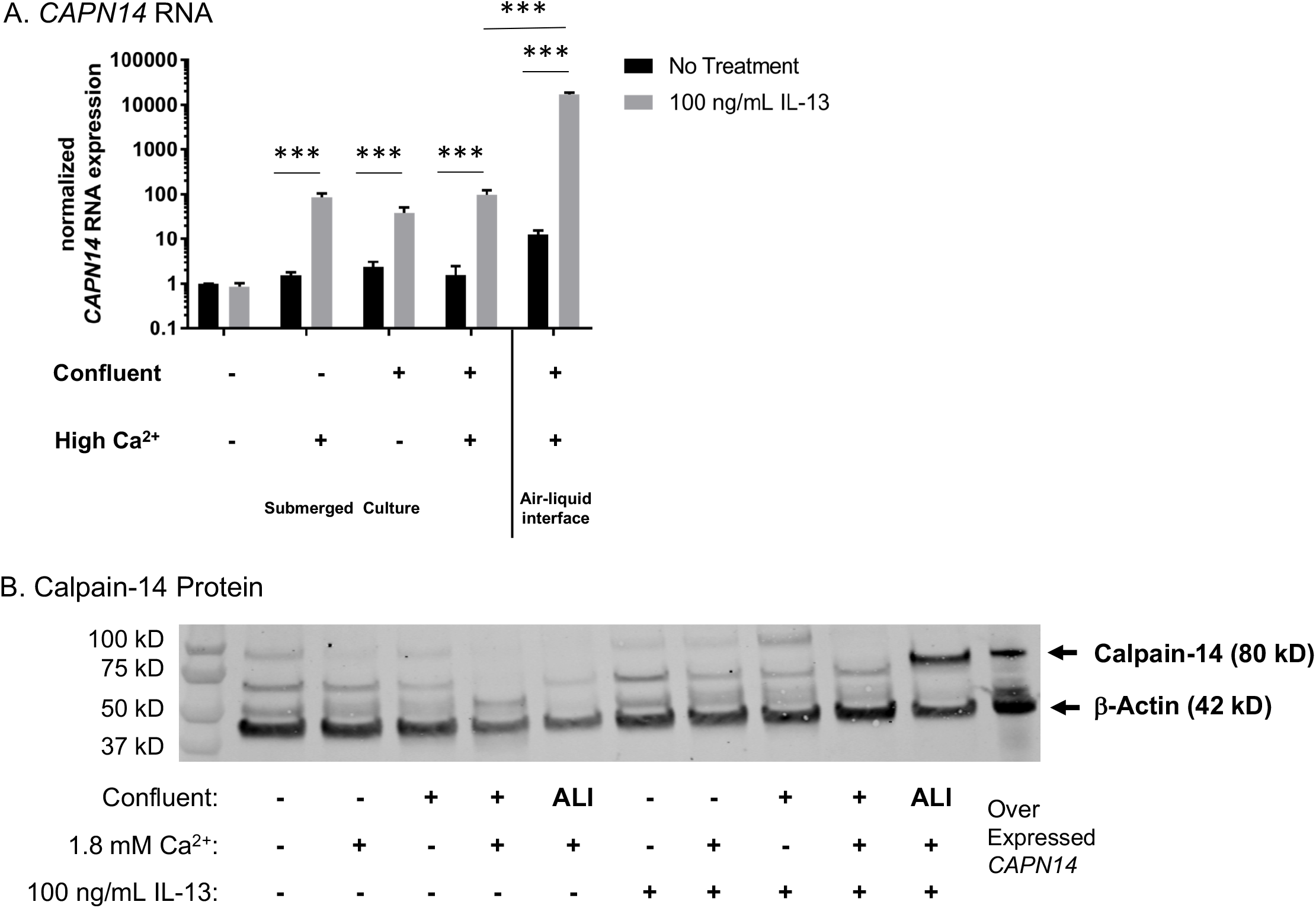
The expression of *CAPN14* in esophageal epithelial cells is dependent upon calcium, confluency, and IL-13. EPC2 esophageal epithelial cells were grown to various levels of confluence (80%-indicated by -, or 100%-indicated by +) with relatively low (0.09 mM) or high (1.8 2+ mM) Ca^2+^ and with or without 100 ng/mL IL-13 for 24 hours. Cultures were grown either as a monolayer submerged culture or in an air-liquid interface (ALI) setup. *A*, RNA was isolated and *CAPN14* was measured relative to the expression of the house keeping gene *GAPDH*. All RNA expression values are normalized to cells at 80% confluence without high calcium or IL-13. *B*, Calpain-14 protein expression levels. EPC2 cells that constitutively express *CAPN14* behind a CMV promoter were used as a positive control for calpain-14 protein expression. RNA and protein level expression: ***, t-test p-value<0.001, n=3-5 per group, data representative of 4 independent experiments.

## DISCUSSION

Altogether, we identified immunological (IL-13/IL-4/STAT6), genetic (rs76562819), and epithelial differentiation factors that regulate the expression of *CAPN14*. These results are important because calpain-14 represents a candidate for treatment and prevention of EoE. Previous studies supported a model in which IL-13 induction of *CAPN14* occurs in a STAT6-dependent manner. We identified the STAT6 binding sites, the necessity of these sites for *CAPN14* promoter activity, and the binding of STAT6 to the promoter of *CAPN14* in esophageal cells stimulated with IL-13. We demonstrated genotype-dependent expression of *CAPN14* and identified rs76562819 as the genetic variant that is sufficient to produce genotype-dependent promoter activity. Indeed, the 40.0% reduction of *CAPN14* promoter activity in the genotype-dependent reporter assays mirrors the effect size of the eQTLs measured in subjects with and without EoE (Kottyan et al. 2014). The reporter experiments did not demonstrate statistically significant genotype-dependent promoter activity in cells that were not stimulated with IL-13. This could be due to several factors, including the sensitivity of the reporter system, the non-differentiated context of the reporter experiments, or the fact that IL-13 is needed for genotype-dependent *CAPN14* promoter activity. The fact that relatively undifferentiated esophageal epithelial cell culture systems are sufficient for luciferase reporter systems could be due to how differentiation affects chromatin availability – a factor that is not assessed in the cytoplasmic luciferase reporter assay. *CAPN14* mRNA and protein expression levels strongly support the conclusion that both differentiation and stimulation with IL-13 or IL-4 is critical for the highest expression of calpain-14 in esophageal epithelial cells. The requirement for esophageal epithelial cell differentiation before *CAPN14* expression might be the primary mechanism through which *CAPN14’s* expression is-induced in the esophageal mucosa.

IL-13 is a critical cytokine in the pathoetiology of EoE (Blanchard et al. 2011; O’Shea et al. 2017; Davis et al. 2016; D’Mello et al. 2016; Cianferoni and Spergel 2016; Rothenberg et al. 2015; Sherrill et al. 2014b; Sherrill et al. 2014a; Zuo et al. 2010; Blanchard et al. 2007; Mishra and Rothenberg 2003). Indeed, treating esophageal epithelial cells with IL-13 is an effective way to model approximately 25% of the transcriptome and cytokine secretion profiles found in the esophageal biopsies of patients with EoE (Sherrill et al. 2014b; Lu et al. 2012; Blanchard et al. 2007). IL-13 directly controls the expression of chemokines that recruit eosinophils to the esophagus *(CCL11, CCL24, CCL26)* through STAT6-dependent mechanisms (Lim et al. 2011). IL-13 treatment can also activate other cellular and transcriptional regulators including the protein kinase B (Akt) and mitogen-activated protein kinase (MAPK) pathways (Demoulin and Renauld 1998; Harris et al. 2007), and transcriptional regulation downstream of IL-13 signaling can also be indirect (e.g. genes transcribed in response to IL-13 can regulate the expression of other genes). It was therefore important to establish that two STAT6 sites in the *CAPN14* promoter control IL-13-stimulated expression *of CAPN14*. IL-13 shares many functional properties with IL-4, stemming from the fact that they share a common receptor subunit, the alpha subunit of the IL-4 receptor (Hershey 2003). Both IL-13 and IL-4 signaling lead to STAT6 dimerization, phosphorylation, and nuclear translocation (Kuperman and Schleimer 2008), so it is therefore not surprising that IL-4 might also regulate *CAPN14* mRNA expression and promoter activity (**Figures S1 and S3**). Despite the evidence presented in this study, future experiments in esophageal epithelial cells deficient in STAT6 signaling are needed to fully conclude that STAT6 is necessary in the transcription of *CAPN14*.

Very little is known about the factors regulating the expression, function, and tissue specificity of *CAPN14* (Litosh et al. 2017; Davis et al. 2016; Sleiman et al. 2014; Ueta et al. 2010; Kottyan et al. 2014; Dear and Boehm 2001). A growing body of literature suggests that there are factors specific to the esophagus that affect gene regulation (Keane et al. 2015; Rochman et al. 2018; Rochman et al. 2017). These and other studies are consistent with our findings that while some genes regulated by IL-13 are expressed in various epithelial context (e.g. CCL26/eotain-3), other genes have more nuainced regulatory factors affecting expression (e.g. CAPN14/calpain-14). In this study, we demonstrated that epithelial differentiation was important; however, we were unable to establish specific tissue-specific molecular pathways that lead to the differences in CCL26 and CAPN14 expression in response to IL-13.

This study focused on the regulation of *CAPN14* transcription. Future studies will assess the post-translational regulation and activity of calpain-14. The structure of calpain-14 strongly suggests that it might dimerize with calpain-S1 (Litosh et al. 2017). Other members of the calpain family are known to undergo autolysis (Ono et al. 2016a; Ermolova et al. 2011; Taveau et al. 2003; Badugu et al. 2008), and proteomic analysis of calpain-14 expression found evidence consistent with cleavage of calpain-14, especially in non-differentiated conditions (**Figure 4 B**). We used an overexpression construct of calpain-14 to identify the specific band bound by the anti-calpain-14 antibody in the Western blots (**Figure 4 B**). Calpains cleave substrates in a calcium-dependent manner based upon the conformation, rather than the amino acid sequence, of their targets (Ono et al. 2016b; Sorimachi et al. 2012). Full biochemical analyses of the substrates of calpain-14 will be critical as the functions of calpain-14 in health and EoE are discovered.

The current study identifies critical factors that regulate *CAPN14* expression. Altogether, this study makes an important advance towards identification of the factors that control the genotype-dependent transcriptional regulation of *CAPN14* in esophageal epithelium.

## ACKNOWLEDGMENTS

This work was supported by National Institutes of Health [R37 AI045898, R01 AI124355, R01 DK107502, R01 NS099068, R21 HG008186, P30 AR070549, U01 HG008666]; the Campaign Urging Research for Eosinophilic Disease (CURED); the Buckeye Foundation; the Sunshine Charitable Foundation and its supporters, Denise A. Bunning and David G. Bunning; the American Partnership of Eosinophilic Disorders (APFED); Cincinnati Children’s Hospital CpG Pilot Study and Endowed Scholar awards to M.T.W.

This project was supported in part by PHS Grant P30 DK078392 DNA sequencing core of the Digestive Disease Research Core Center in Cincinnati. N.W. and S.A.C. were supported by NIH Grant DA038017 and funds from CCRF, including an Arnold Strauss Fellowship to S.A.C.

## REFERENCES

Alexander, E.S., L.J. Martin, M.H. Collins, L.C. Kottyan, H. Sucharew et al., 2014 Twin and family studies reveal strong environmental and weaker genetic cues explaining heritability of eosinophilic esophagitis. J Allergy Clin Immunol 134 (5):1084–1092 e1081.

Badugu, R., M. Garcia, V. Bondada, A. Joshi, and J.W. Geddes, 2008 N terminus of calpain 1 is a mitochondrial targeting sequence. J Biol Chem 283 (6):3409–3417.

Blanchard, C., M.K. Mingler, M. Vicario, J.P. Abonia, Y.Y. Wu et al., 2007 IL-13 involvement in eosinophilic esophagitis: transcriptome analysis and reversibility with glucocorticoids. J Allergy Clin Immunol 120 (6):1292–1300.

Blanchard, C., E.M. Stucke, B. Rodriguez-Jimenez, K. Burwinkel, M.H. Collins et al., 2011 A striking local esophageal cytokine expression profile in eosinophilic esophagitis. J Allergy Clin Immunol 127 (1):208–217, 217 e201–207.

Blanchard, C., N. Wang, K.F. Stringer, A. Mishra, P.C. Fulkerson et al., 2006 Eotaxin-3 and a uniquely conserved gene-expression profile in eosinophilic esophagitis. J Clin Invest 116 (2):536–547.

Bray, N.L., H. Pimentel, P. Melsted, and L. Pachter, 2016 Near-optimal probabilistic RNA-seq quantification. Nat Biotechnol 34 (5):525–527.

Cianferoni, A., and J. Spergel, 2016 Eosinophilic Esophagitis: A Comprehensive Review. Clin Rev Allergy Immunol 50 (2):159–174.

D’Mello, R.J., J.M. Caldwell, N.P. Azouz, T. Wen, J.D. Sherrill et al., 2016 LRRC31 is induced by IL-13 and regulates kallikrein expression and barrier function in the esophageal epithelium. Mucosal Immunol 9 (3):744–756.

Davis, B.P., E.M. Stucke, M.E. Khorki, V.A. Litosh, J.K. Rymer et al., 2016 Eosinophilic esophagitis-linked calpain 14 is an IL-13-induced protease that mediates esophageal epithelial barrier impairment. JCI Insight 1 (4):e86355.

Dear, T.N., and T. Boehm, 2001 Identification and characterization of two novel calpain large subunit genes. Gene 274 (1–2):245–252.

Demoulin, J.B., and J.C. Renauld, 1998 Signalling by cytokines interacting with the interleukin-2 receptor gamma chain. Cytokines Cell Mol Ther 4 (4):243–256.

Ermolova, N., E. Kudryashova, M. DiFranco, J. Vergara, I. Kramerova et al., 2011 Pathogenity of some limb girdle muscular dystrophy mutations can result from reduced anchorage to myofibrils and altered stability of calpain 3. Hum Mol Genet 20 (17):3331–3345.

Ferreira, M.A., M.C. Matheson, D.L. Duffy, G.B. Marks, J. Hui et al., 2011 Identification of IL6R and chromosome 11q13.5 as risk loci for asthma. Lancet 378 (9795):1006–1014.

Ferreira, M.A., M.C. Matheson, C.S. Tang, R. Granell, W. Ang et al., 2014 Genome-wide association analysis identifies 11 risk variants associated with the asthma with hay fever phenotype. J Allergy Clin Immunol 133 (6):1564–1571.

Flicek, P., I. Ahmed, M.R. Amode, D. Barrell, K. Beal et al., 2013 Ensembl 2013. Nucleic Acids Res 41 (Database issue):D48–55.

Greisenegger, E.K., F. Zimprich, A. Zimprich, A. Gleiss, and T. Kopp, 2013 Association of the chromosome 11q13.5 variant with atopic dermatitis in Austrian patients. Eur J Dermatol 23 (2):142–145.

Harris, J., S.A. De Haro, S.S. Master, J. Keane, E.A. Roberts et al., 2007 T helper 2 cytokines inhibit autophagic control of intracellular Mycobacterium tuberculosis. Immunity 27 (3):505–517.

Hershey, G.K., 2003 IL-13 receptors and signaling pathways: an evolving web. J Allergy Clin Immunol 111 (4):677–690; quiz 691.

Hinds, D.A., G. McMahon, A.K. Kiefer, C.B. Do, N. Eriksson et al., 2013 A genome-wide association meta-analysis of self-reported allergy identifies shared and allergy-specific susceptibility loci. Nat Genet 45 (8):907–911.

Kartashov, A.V., and A. Barski, 2015 BioWardrobe: an integrated platform for analysis of epigenomics and transcriptomics data. Genome Biol 16:158.

Kc, K., M.E. Rothenberg, and J.D. Sherrill, 2015 In vitro model for studying esophageal epithelial differentiation and allergic inflammatory responses identifies keratin involvement in eosinophilic esophagitis. PLoS One 10 (6):e0127755.

Keane, T.J., A. DeWard, R. Londono, L.T. Saldin, A.A. Castleton et al., 2015 Tissue-Specific Effects of Esophageal Extracellular Matrix. Tissue Eng Part A 21 (17–18):2293–2300.

Kottyan, L.C., B.P. Davis, J.D. Sherrill, K. Liu, M. Rochman et al., 2014 Genome-wide association analysis of eosinophilic esophagitis provides insight into the tissue specificity of this allergic disease. Nat Genet 46 (8):895–900.

Kottyan, L.C., M.T. Weirauch, and M.E. Rothenberg, 2015 Making it big in allergy. J Allergy Clin Immunol 135 (1):43–45.

Kuperman, D.A., and R.P. Schleimer, 2008 Interleukin-4, interleukin-13, signal transducer and activator of transcription factor 6, and allergic asthma. Curr Mol Med 8 (5):384–392.

Li, J., Y. Zhang, and L. Zhang, 2015 Discovering susceptibility genes for allergic rhinitis and allergy using a genome-wide association study strategy. Curr Opin Allergy Clin Immunol 15 (1):33–40.

Liacouras, C.A., G.T. Furuta, I. Hirano, D. Atkins, S.E. Attwood et al., 2011 Eosinophilic esophagitis: updated consensus recommendations for children and adults. J Allergy Clin Immunol 128 (1):3–20 e26; quiz 21-22.

Lim, E.J., T.X. Lu, C. Blanchard, and M.E. Rothenberg, 2011 Epigenetic regulation of the IL-13-induced human eotaxin-3 gene by CREB-binding protein-mediated histone 3 acetylation. J Biol Chem 286 (15):13193–13204.

Litosh, V.A., M. Rochman, J.K. Rymer, A. Porollo, L.C. Kottyan et al., 2017 Calpain-14 and its association with eosinophilic esophagitis. J Allergy Clin Immunol 139 (6):1762–1771 e1767.

Livak, K.J., and T.D. Schmittgen, 2001 Analysis of relative gene expression data using real-time quantitative PCR and the 2(-Delta Delta C(T)) Method. Methods 25 (4):402–408.

Lu, T.X., E.J. Lim, T. Wen, A.J. Plassard, S.P. Hogan et al., 2012 MiR-375 is downregulated in epithelial cells after IL-13 stimulation and regulates an IL-13-induced epithelial transcriptome. Mucosal Immunol 5 (4):388–396.

Macara, I.G., R. Guyer, G. Richardson, Y. Huo, and S.M. Ahmed, 2014 Epithelial homeostasis. Curr Biol 24 (17):R815–825.

Mishra, A., and M.E. Rothenberg, 2003 Intratracheal IL-13 induces eosinophilic esophagitis by an IL-5, eotaxin-1, and STAT6-dependent mechanism. Gastroenterology 125 (5):1419–1427.

Namjou, B., K. Marsolo, R.J. Caroll, J.C. Denny, M.D. Ritchie et al., 2014 Phenome-wide association study (PheWAS) in EMR-linked pediatric cohorts, genetically links PLCL1 to speech language development and IL5-IL13 to Eosinophilic Esophagitis. Front Genet 5:401.

O’Shea, K.M., S.S. Aceves, E.S. Dellon, S.K. Gupta, J.M. Spergel et al., 2017 Pathophysiology of Eosinophilic Esophagitis. Gastroenterology.

Ono, Y., K. Ojima, F. Shinkai-Ouchi, S. Hata, and H. Sorimachi, 2016a An eccentric calpain, CAPN3/p94/calpain-3. Biochimie 122:169–187.

Ono, Y., T.C. Saido, and H. Sorimachi, 2016b Calpain research for drug discovery: challenges and potential. Nat Rev Drug Discov 15 (12):854–876.

Ramasamy, A., I. Curjuric, L.J. Coin, A. Kumar, W.L. McArdle et al., 2011 A genome-wide meta-analysis of genetic variants associated with allergic rhinitis and grass sensitization and their interaction with birth order. J Allergy Clin Immunol 128 (5):996–1005.

Rochman, M., N.P. Azouz, and M.E. Rothenberg, 2018 Epithelial origin of eosinophilic esophagitis. J Allergy Clin Immunol 142 (1):10–23.

Rochman, M., A.V. Kartashov, J.M. Caldwell, M.H. Collins, E.M. Stucke et al., 2015 Neurotrophic tyrosine kinase receptor 1 is a direct transcriptional and epigenetic target of IL-13 involved in allergic inflammation. Mucosal Immunol 8 (4):785–798.

Rochman, M., J. Travers, J.P. Abonia, J.M. Caldwell, and M.E. Rothenberg, 2017 Synaptopodin is upregulated by IL-13 in eosinophilic esophagitis and regulates esophageal epithelial cell motility and barrier integrity. JCI Insight 2 (20).

Rothenberg, M.E., T. Wen, A. Greenberg, O. Alpan, B. Enav et al., 2015 Intravenous anti-IL-13 mAb QAX576 for the treatment of eosinophilic esophagitis. J Allergy Clin Immunol 135 (2):500–507.

Sherrill, J.D., K. Kc, D. Wu, Z. Djukic, J.M. Caldwell et al., 2014a Desmoglein-1 regulates esophageal epithelial barrier function and immune responses in eosinophilic esophagitis. Mucosal Immunol 7 (3):718–729.

Sherrill, J.D., K.C. Kiran, C. Blanchard, E.M. Stucke, K.A. Kemme et al., 2014b Analysis and expansion of the eosinophilic esophagitis transcriptome by RNA sequencing. Genes Immun 15 (6):361–369.

Sherrill, J.D., and M.E. Rothenberg, 2014 Genetic and epigenetic underpinnings of eosinophilic esophagitis. Gastroenterol Clin North Am 43 (2):269–280.

Sleiman, P.M., M.L. Wang, A. Cianferoni, S. Aceves, N. Gonsalves et al., 2014 GWAS identifies four novel eosinophilic esophagitis loci. Nat Commun 5:5593.

Sorimachi, H., S. Hata, and Y. Ono, 2011 Impact of genetic insights into calpain biology. J Biochem 150 (1):23–37.

Sorimachi, H., H. Mamitsuka, and Y. Ono, 2012 Understanding the substrate specificity of conventional calpains. Biol Chem 393 (9):853–871.

Tang, X.F., H.Y. Tang, L.D. Sun, F.L. Xiao, Z. Zhang et al., 2012 Genetic variant rs4982958 at 14q11.2 is associated with allergic rhinitis in a Chinese Han population running title: 14q11.2 is a susceptibility locus for allergic rhinitis. J Investig Allergol Clin Immunol 22 (1):55–62.

Taveau, M., N. Bourg, G. Sillon, C. Roudaut, M. Bartoli et al., 2003 Calpain 3 is activated through autolysis within the active site and lyses sarcomeric and sarcolemmal components. Mol Cell Biol 23 (24):9127–9135.

Trapnell, C., L. Pachter, and S.L. Salzberg, 2009 TopHat: discovering splice junctions with RNA-Seq. Bioinformatics 25 (9):1105–1111.

Ueta, M., K. Mizushima, N. Yokoi, Y. Naito, and S. Kinoshita, 2010 Expression of the interleukin-4 receptor alpha in human conjunctival epithelial cells. Br J Ophthalmol 94 (9):1239–1243.

Vallabh, S., A.V. Kartashov, and A. Barski, 2018 Analysis of ChIP-Seq and RNA-Seq Data with BioWardrobe. Methods Mol Biol 1783:343–360.

Weidinger, S., S.A. Willis-Owen, Y. Kamatani, H. Baurecht, N. Morar et al., 2013 A genome-wide association study of atopic dermatitis identifies loci with overlapping effects on asthma and psoriasis. Hum Mol Genet 22 (23):4841–4856.

Wilming, L.G., J.G. Gilbert, K. Howe, S. Trevanion, T. Hubbard et al., 2008 The vertebrate genome annotation (Vega) database. Nucleic Acids Res 36 (Database issue):D753–760.

Zuo, L., P.C. Fulkerson, F.D. Finkelman, M. Mingler, C.A. Fischetti et al., 2010 IL-13 induces esophageal remodeling and gene expression by an eosinophil-independent, IL-13R alpha 2-inhibited pathway. J Immunol 185 (1):660–669.

